# A novel positive feedback mechanism of ABI5 phosphorylation by mitogen activated protein kinase-3 regulates ABA signaling in *Arabidopsis*

**DOI:** 10.1101/2021.02.02.429361

**Authors:** Prakash Kumar Bhagat, Deepanjali Verma, Neetu Verma, Alok Krishna Sinha

**Affiliations:** National Institute of Plant Genome Research, Aruna Asaf Ali Marg, New Delhi, 110067, India

**Keywords:** ABA, AtABI5, AtMPK3, AtMPK6, Drought stress, Dimerization, Floral transition, MAP kinase, Phosphorylation, Post-germination growth, seed germination

## Abstract

Seed germination is the crucial physiological process regulated by both environmental and endogenous phytohormones. ABA negatively regulates seed germination, post-germination growth and floral transition, however, the cross talk between multiple regulatory pathways are still unclear. Here, we show that ABA activates two MAP kinases, AtMPK3/AtMPK6 and selectively regulates the transcript of AtMPK3 through ABI5, a master regulator of ABA signaling. As a feedback loop, AtMPK3 interacts and phosphorylates ABI5 at the serine-314 position. ABI5 phosphorylation by MAP kinases positively regulates ABI5 nuclear localization and negatively regulates its dimerization. Subcellular localization of ABI5 phospho-null protein further suggests the role of phosphorylation in regulation of its cytoplasmic stability and its nuclear dimerization. Overexpression of phosphor-null ABI5 in *abi5-8* mutant restored the ABA sensitivity during seed germination and delayed the floral transition as compared to phospho-mimic ABI5. Additionally, overexpression of constitutive phosphorylated ABI5 in *abi5-8* mutants suggest that phosphorylation makes ABI5 partially inactive. Furthermore, phospho-null ABI5 plants showed drought sensitive phenotype whereas, *mpk3*, *mkk4*, *mkk5*, *abi5-8* and phosphor-mimic plants showed drought tolerant phenotype. Our findings present a new insight between MAP kinase cascade and ABA signaling which collectively regulates the ABA response through ABI5 phosphorylation.

## Introduction

During abiotic stress, plants employ a conserved abscisic acid (ABA) signaling which integrates multiple complex pathways that are regulated both at transcriptional and post-translational levels (Nakashima et al., 2009). A well-studied ABA pathway in Arabidopsis involves an ABA receptor and a core ABA signaling kinase-phosphatase modules (Fujii et al., 2007). Activated kinase phosphorylates ABA-responsive transcription factors (TFs) such as ABF and a bZIP transcription factor, ABSCISIC ACID-INSENSITIVE 5 (ABI5) (Nakamura et al., 2001; Nakashima et al., 2009; Dai et al., 2013). ABA signaling regulates the plant growth and development from seed germination to that of floral transition and also integrates the stress responses involving cross talk with multiple signaling and hormone pathways (Piskurewicz et al., 2008; Wang et al., 2013).

ABI5 is a master regulator of ABA signaling and knockout of *ABI5* results in ABA-insensitive phenotype which was used to characterise a number of new factors in ABA metabolism and signaling (Lopez-Molina and Chua, 2000; Lopez-Molina et al., 2002). Presence of ABA strictly regulates the expression and function of ABI5 at both transcriptional and post-translational stages by covalent modifications (Miura et al., 2009; Nakashima et al., 2009; Liu and Stone, 2010, 2013). Transgenic plants overexpressing ABI5 are hypersensitive to ABA, however they grow normally in the absence of ABA. This suggests that ABA induced post-translational modifications (PTMs) are crucial for ABA–induced regulations (Lopez-Molina et al., 2001). Additionally, floral transition is early in the *abi5* mutants and its overexpression delays flowering. Thus, ABA exerts the inhibitory effect on floral transition through ABI5 which upregulates the expression of *FLOWERING LOCUS C* (*AtFLC*) which in turn inhibits the flowering (Wang et al., 2013). However, floral transition is also regulated by other factors such as ABI4, independent of ABI5 (Shu et al., 2018). Therefore, floral transition is a complex mechanism governed by ABA.

Mitogen-activated protein kinases (MAPK) pathway is an evolutionarily conserved signaling cascade which regulates several developmental and stress responses (Jalmi et al., 2018). During abiotic or drought stress, the ABA content is significantly enhanced and is known to activate several MAP kinases (Matsuoka et al., 2015; Li et al., 2017b). Role of MAPK cascades involved in plant growth and development as well as in stress responses have been elucidated (Xu and Zhang, 2015; Jalmi et al., 2018; Verma et al., 2019). The link between ABA and MAP kinase cascade is specifically well explored in primary root growth, stomatal responses and leaf senescence (Danquah et al., 2015; Khokon et al., 2015; Matsuoka et al., 2015; Li et al., 2017a). In the guard cells, ROS activates MPK9 and MPK12 which regulates ABA response positively (Jammes et al., 2009). It has been reported during ABA signaling MAP kinase signaling transduces the signal to the transcriptional apparatus (Lu et al., 2002). Though, significant contributions have been made in understanding the role of MAP kinase cascade and ABA signaling during stomatal response which is usually a late growth response, the information on early response on seed germination and post germination growth is still missing. The underlying mechanism for this crosstalk at molecular level needs further investigation.

We here report a novel regulation of AtABI5 by AtMPK3 and their role in seed germination, flowering and drought tolerance. We identify activation of two MAP kinases, AtMPK3 and AtMPK6 during exogenous application of ABA. Interestingly, ABA transcriptionally upregulates AtMPK3 through AtABI5 by binding to the ABA-responsive element present selectively in AtMPK3 but absent in AtMPK6. Additionally, the Activated AtMPK3 interacts with AtABI5 in the nucleus and phosphorylates it at an evolutionarily conserved serine-314 residue. Phosphorylation of ABI5 regulates its subcellular localization and inhibits its dimerization. In addition, we further show that only phospho-null ABI5 could restore the ABA sensitivity when overexpressed in *abi5-8* mutant background. However, both phosphor-null and phosphor-mimic ABI5 transgenic plants exhibited delayed flowering but this phenotype was more pronounced in phospho-null transgenic plants. Drought stress analysis of MAP kinase mutants and transgenic plants suggested that except phosphor-null ABI5, all other mutants showed drought tolerance phenotype and better recovery.

## Materials and Methods

### Plant materials and growth conditions

The WT (Col-0) and mutant seeds were grown as described in Verma et. al., 2019. The MS plates were supplemented with indicated concentrations of ABA (Sigma) for seed germination and ABA sensitivity assay. For production of overexpression transgenic plants, the coding sequences of ABI5 and its mutant variants (ABI5^S314A^ and ABI5^S314D^) were cloned in pGWB5 binary vector using Gateway cloning method and transformed into *Agrobacterium tumefaciens* GV3101 strain. Positive constructs were introduced into *abi5-8* mutant plants by floral-dip method. The subsequent seeds were screened on ½ MS plates supplemented with kanamycin and hygromycin. T3 generation seeds were used for further experiments.

### Plant protein extraction and Immunoblotting

Arabidopsis seedlings were ground into fine powder using liquid nitrogen. Total proteins were extracted using extraction buffer (50 mM HEPES, pH 7.5, 5 mM EDTA, 5 mM EGTA, 10 mM DTT, 10 mM Na_3_VO_4_, 10 mM NaF, 50 mM β-glycerolphosphate, 1 mM phenylmethylsulfonyl fluoride, protease inhibitor cocktail, and 10% glycerol). Proteins were denatured by boiling with a 5X sodium dodecyl sulfate (SDS) loading buffer for 3 minutes. Immunoblotting was performed with phospho-ERK1/2 (pERK1/2) antibody (Cell signaling), anti-AtMPK3 (Sigma) and anti-AtMPK6 (Sigma) as described earlier (Singh and Sinha, 2016).

### ABA sensitivity assay

WT and mutant seeds of *mpk3* and *abi5* were plated on ½ MS agar plate either without ABA (control) or with ABA (1μM, 2 μM, 3 μM and 5 μM). After vernalisation for 2 to 3 days, plates were transferred to constant light for germination. Germinated seeds were captured using a Nikon stereo microscope.

### RNA isolation and quantitative real time PCR (qRT-PCR) analysis

Total RNA was extracted from control and treated samples using Trizol reagent sigma (#T9424). The cDNA was prepared using RevertAid H Minus First Strand cDNA synthesis kit (#K1632, Thermo Scientific) and qRT-PCR were performed as described previously (Singh and Sinha, 2016). The primer sets used in the analysis are given in the Table S1.

### Yeast-two hybrid assay

The interaction between MPK3 and ABI5 was checked using the Matchmaker yeast two-hybrid system (BD Biosciences) as described previously with slight modifications (Sheikh et al., 2013; Raghuram et al., 2015). Briefly, the N-/C-terminal half of ABI5 along with its full length CDS and full length MPK3 were cloned in the pGADT7 and pGBKT7 vectors. The yeast transformation was performed according to the Fast yeast transformation kit (Gbioscience: GZ‐1).

### *In-vitro* GST-pulldown assay

For in-vitro protein-protein interaction study, bacterially purified GST-AtMPK3 and His-AtABI5 were incubated in the binding buffer (50 mm Tris-HCl, pH 7.5, 150 mm NaCl, 0.2% (v/v) glycerol, 0.1% (v/v) Triton X-100, 1 mm EDTA, pH 8.0, 1 mm PMSF, 0.1% (v/v) Nonidet P-40) over night at 4°C. After washing with 1x PBS for three-four times, the bound proteins were separated on 10% SDS-PAGE. Interactor proteins were detected by immunoblot with anti-his and anti-AtMPK3 antibodies.

### Bimolecular fluorescence complementation (BiFC) assay and sub-cellular localization

For BiFC assay, MPK3 and ABI5 (along with mutant ABI5^S314A^ and ABI5^S314D^) were cloned in frame with binary vectors pSPYCE M and pSPYNE 173 which contains C- terminal half (cYFP) and n-terminal half (nYFP) of yellow fluorescence protein (YFP), respectively. For localization studies, the ABI5 WT and mutant variants (ABI5^S314A^and ABI5^S314D^) were cloned in pGWB5 which has green fluorescence protein (GFP) at its amino- terminal. *Nicotiana benthamiana* leaves were infiltrated and fluorescence were monitored as described previously (Raghuram et al., 2015).

### Bacterial protein expression and in-vitro phosphorylation assay

MPK3 and ABI5 CDS were cloned in frame with GST (pGEX4t2) and 6xHis (pET28a) tags, respectively. The proteins were induced in BL21 strain of *E. coli* by addition of 1mM IPTG at 28°C. The recombinant proteins tagged with GST and 6xHis were purified by affinity chromatography using GST-sepharose beads and Ni-NTA beads, respectively. The total proteins were quantified by Bradford reagent (Sigma) and purity was analysed by SDS-PAGE. The *in-vitro* phosphorylation of ABI5 was performed as described previously (Raghuram et al., 2015).

### Multiple protein sequence alignment and Site - directed mutagenesis (SDM) analysis

The multiple protein sequences were aligned from different plant species with *Arabidopsis thaliana* ABI5 and the conserved amino acids were analysed using the Uniprot database (https://www.uniprot.org/). The putative and invariant amino acids were marked and indicated. The conserved invariant and single putative MAP kinase site, S-314 was used for SDM analysis. The list of primers used is mentioned in Table S1, primers list. The ABI5-pET28a and ABI5-pENTR vectors were used as templates for PCR based SDM. The PCR reaction was digested with DpnI restriction enzyme overnight and the resultant reaction was transformed in *E. coli* DH5α competent cells. The mutation was confirmed by DNA sequencing.

### Electrophoretic mobility shift assay (EMSA)

The DNA-protein interaction between AtMPK3 promoter and ABI5 transcription factor was performed by EMSA. The DNA sequence containing the ACGT core element (−402 to −256 base pair) were PCR amplified, radiolabelled and used for the binding experiment with bacterially purified ABI5-His proteins as described previously (Singh and Sinha, 2016). Briefly, radiolabelled probes were either incubated alone or with increasing protein concentration of His-ABI5 in the reaction buffer (20mM HEPES, pH 7.4; 10mM KCl; 0.5mM EDTA; 0.5mM DTT; 1mM MgCl_2_; 3% Glycerol and 1 μg of poly(dI-dC) at room temperature for 30 minutes. The DNA-protein complex was separated on 4% PAGE using 0.5x TBE running buffer and auto-radiographed using typhoon.

### Flowering phenotype analysis

The flowering phenotype was analysed according to the previous study (Wang et al., 2013; Shu et al., 2018). Seeds were directly sown on the potted soil and flowering phenotypes were monitored till bolting. The images were taken every day.

### Drought stress analysis

The drought stress response was conducted using *mpk3*, *mkk4*, *mkk5*, *abi5-8*, *ABI5*^*S314A*^*::abi5-8* and *ABI5*^*S314D*^*::abi5-8* transgenic plants according to previously described (Harb et al., 2010). The soil grown 25 to 30 days old plants were subjected to drought stress conditions by further growing the plants without water until the plants showed drying and leaf chlorosis phenotype. To examine the recovery response, plants were re-watered and successively recovery was monitored every day. The plants which were unable to recover, considered as sensitive and those recovered with green leaves, termed as tolerant.

## Results

### ABA regulates activation, transcription and translation of MPK3

Abiotic stresses or exogenous ABA application often activate MAP kinase cascade (Danquah et al., 2015). To further elucidate the crosstalk between MAP kinases and ABA signaling we first analysed the activation of MAP kinase under increasing concentration of exogenous ABA application. The activation of two MAP kinases at 42 and 46 kDa molecular weight was observed **(Figure 1A)**. We then monitored the protein level of two MAP kinases corresponding to the 42 and 46 kDa of AtMPK3 and AtMPK6, respectively **(Figure 1A)**. Interestingly, it was found that the protein level of AtMPK3 was upregulated by increasing concentrations of ABA as compared to AtMPK6. To confirm this observation, we performed the same experiment in *mpk3* mutant and probed it with anti-AtMPK3 and anti-AtMPK6 antibodies **(Figure 1B)**. It was observed that AtMPK3 protein increased with time during exogenous ABA application. These results suggested that ABA signalling might regulate the MAP kinase cascade possibly through MAP kinase activation and also by increasing the protein content of AtMPK3. To check whether AtMPK3 gene expression is subjected to transcription regulation by ABA, we analysed the transcript abundance of *AtMPK3* and *AtMPK6*. We found that only the transcript of *AtMPK3* upregulated by ABA **(Figure 1C)**, very much in the same lines to that of its protein abundance observation. Taken together, these data indicate that MAP kinase signaling is regulated by ABA not only at the activation of its protein activity but also at the levels of protein and gene expression of *AtMPK3*.

**Figure 1.**
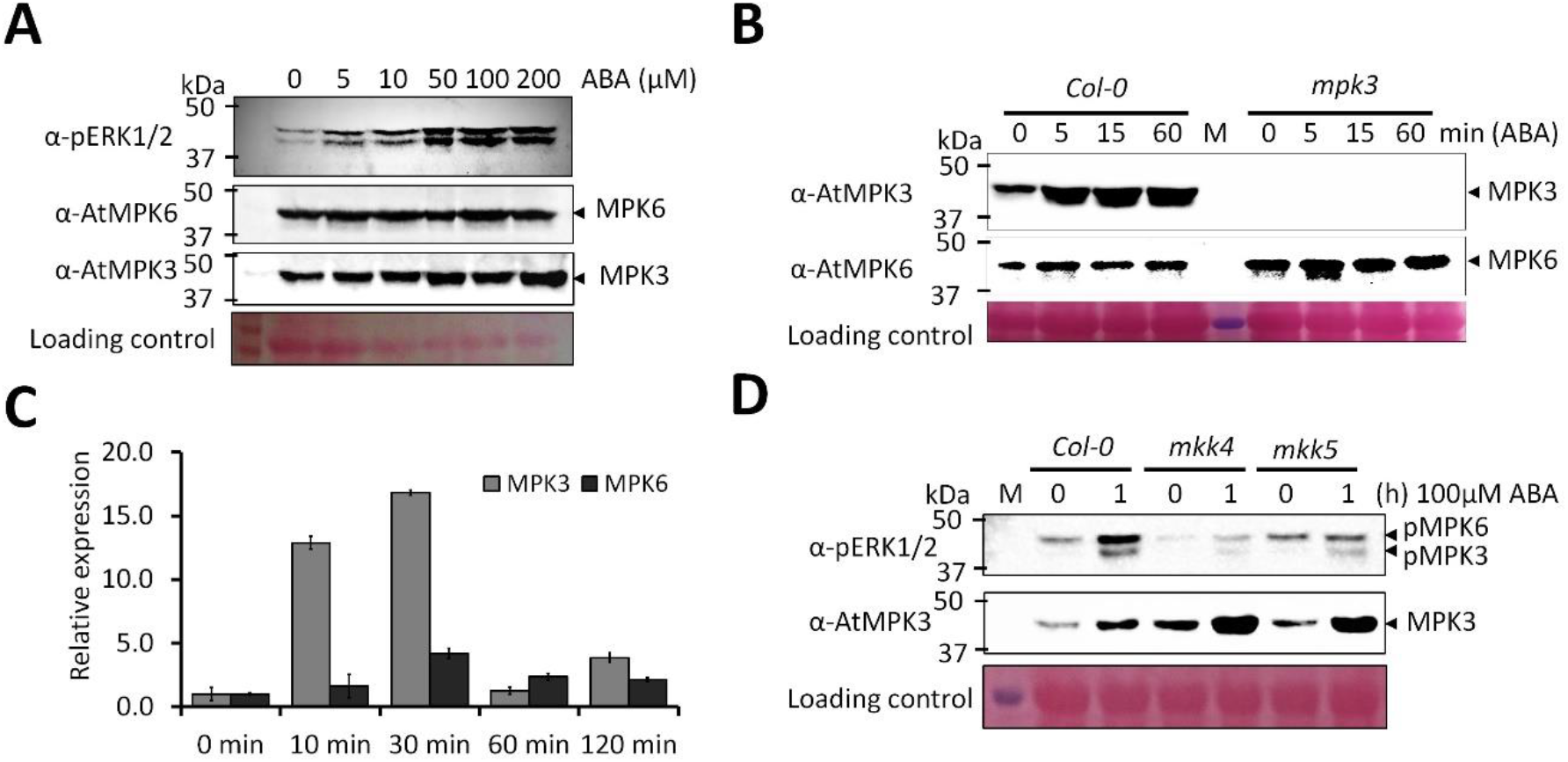
MAP kinase signaling is tightly regulated by ABA. **(A)** Activation of MAP kinases, AtMPK3 and AtMPK5 in wild type plants by ABA treatment. Total protein was isolated from treated plants with indicated concentration of ABA for 30 minutes and activation of MAP kinases were detected by immunoblot analysis using anti-phospho-ERK1/2 (pERK1/2) antibody (top). The protein content of AtMPK6 (middle) and AtMPK6 (bottom) were detected by immunoblot using anti-AtMPK6 and anti-AtMPK3 antibody, respectively. The ponceau staining of rubisco was used as loading control. **(B)** Protein level of AtMPK3 in wild type and *mpk3* mutants under ABA treatment. Seedlings were treated with 100μM ABA for indicated times and total protein was used for immunoblot analysis using anti-AtMPK3 antibody. **(C)** Effect of ABA on AtMPK3 and AtMPK6 transcript. Total RNA was isolated from wild type seedlings treated with 100μM ABA for indicated times. The expression of AtMPK3 and AtMPK6 were determined by quantitative-real time PCR. Data are presented of mean ± s.d. of similar biological replicates. **(D)** AtMKK4 and AtMKK5 are upstream kinases in ABA signaling. The wild type, mkk4 and mkk5 seedlings were treated with 100μM ABA for indicated times and total protein was isolated. Activation of MAP kinases (top) were detected by anti-pERK1/2 antibody and protein abundance of AtMPK3 (bottom) was detected by immunoblot analysis using anti-AtMPK3 antibody. The ponceau stained Rubisco band was used as loading control.

### MKK4/5 are involved in ABA dependent activation of MPK3

To further explore the upstream MAP kinase kinase of AtMPK3 and AtMPK6 in the ABA pathway, we used two previously well-known interactors, AtMKK4 and AtMKK5. We found that activation of these two MAP kinases were compromised in the *mkk4* and *mkk5* mutants suggesting their role in the AtMPK3 and AtMPK6 activation **(Figure 1D)**. In contrast to lower activation of AtMPK3 the ABA significantly upregulated its protein content as compared to wild type **(Figure 1D)**. These data suggest the role of MKK4 and MKK5 in the activation of MAP kinases during ABA treatment but not in protein accumulation of AtMPK3.

### ABI5 interacts with MPK3 promoter and regulates its transcription

The specific upregulation of AtMPK3 transcript and not of AtMPK6 suggested that AtMPK3 might be regulated by one of the ABA responsive transcription factor/s. Analysis of 1kb promoter sequence of AtMPK3 indicated the presence of six core ABA responsive DNA elements (ACGT) **(Figure 2A).** However, there was no such elements present in the promoter of AtMPK6 **(Supplemental Figure 1)**. The two closest core elements in AtMPK3 promoter were present at −269^th^ and 287^th^ position downstream of ATG while the other four at −377^th^, −382^nd^, −890^th^ and −926^th^ positions. To find the putative ABA responsive TF that might regulate the AtMPK3 expression, we analysed the transcript level of a well-characterised ABA master regulator, ABI5. The expression of ABI5 was upregulated by the ABA. **(Figure 2B).** Next, we checked the protein level of AtMPK3 in the wild type, *mpk3* and *abi5-8* mutants in the presence of ABA. We found that AtMPK3 protein decreased in ABA treated samples in the *abi5* mutant as compared to the wild type **(Figure 2C)**. These data suggest that ABI5 regulates AtMPK3 protein expression during ABA signaling. To further test the interaction of AtABI5 with AtMPK3 promoter, we performed the *in-vitro* DNA binding assay employing electrophoretic mobility shift assay (EMSA) **(Figure 2D)**. We used AtMPK3 promoter fragments (−402 to −256) containing four ACGT elements and performed EMSA using HIS-ABI5 protein. We found that increasing ABI5 protein concentrations showed a gel shift and use of unlabelled probe diminished the complex formation, suggesting the specificity of DNA-protein interaction. Taken together these results suggest that ABI5 interacts with AtMPK3 promoter and regulates its expression in an ABA-dependent manner.

**Figure 2.**
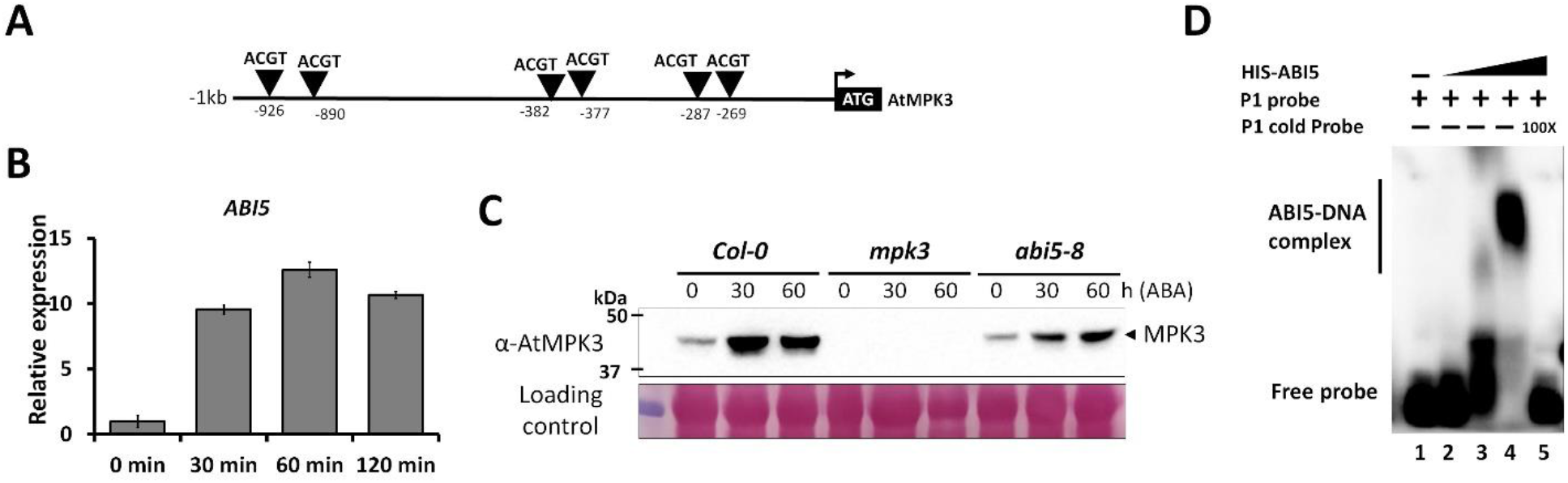
ABA regulates *AtMPK3* expression through *ABI5*. **(A)** Schematic representation of AtMPK3 promoter. −1kb DNA sequence of AtMPK3 promoter was analysed for presence of ABA responsive core element (ABRE) ‘ACGT’. The numbers indicate the nucleotides positions upstream of the transcription start codon, ATG. The black box represents two ABREs and its adjacent nucleotide sequences. **(B)** AtABI5 transcript abundance during ABA treatment. qRT-PCR analysis of AtABI5 transcript in wild type seedlings treated with 100μM ABA. **(C)** Protein abundance of AtMPK3 in *abi5-8* mutant under ABA treatment. Total protein was isolated from wild type, *mpk3* and *abi5-8* treated with 100μM ABA and immunoblot analysis was performed with anti-AtMPK3 antibody.

### ABI5 shows molecular interaction with MPK3

The activation of two MAP kinases, AtMPK3 and AtMPK6 by ABA treatment suggest that MAP kinase cascade might regulate the post-translational modification of ABA responsive transcription factors. To test this possibility, whether ABI5 is an interacting partner of AtMPK3, we first performed the yeast-two hybrid assay using ABI5 variants, full length (ABI5 FL), N-terminal (ABI5N) and C-terminal (ABI5C) **(Figure 3A)**. We found that AtMPK3 strongly interacted with AtABI5 full length and N-terminal but not with the C-terminal variant **(Figure 3B)**. AtABI5-AtMPK3 protein-protein interaction was further validated by *in-vitro* pulldown assay **(Supplemental Figure 2)**. The GST-MPK3 and HIS-AtABI5 were expressed in bacteria, purified and pulled down using GST sepharose beads. The proteins were detected using anti-HIS and anti-AtMPK3 antibodies. The data clearly shows that AtMPK3 and AtABI5 interacts *in-vitro*. To get further insight into the interaction of two proteins *in-vivo*, we performed BiFC assay in tobacco leaves. The interaction between AtABI5 and AtMPK3 was observed within the nucleus **(Figure 3C).** ABI5 is known to regulate the transcription of ABA responsive genes. However, no fluorescence signal was observed in the AtMPK3 and AtABI5 when used independently. Thus, these data suggest that AtABI5 interacts with AtMPK3 *in-planta* and that the interaction takes place in the nucleus.

**Figure 3.**
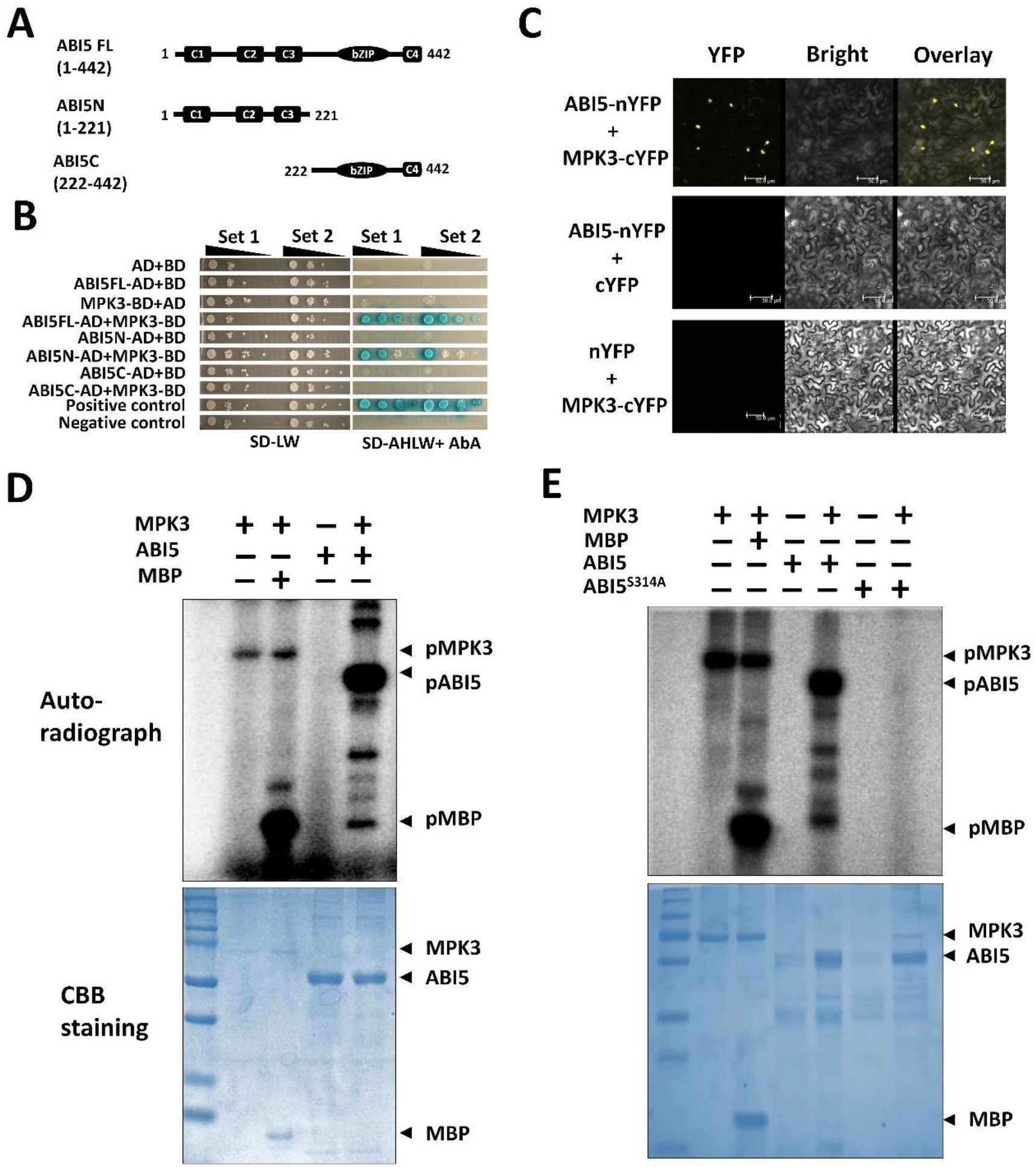
ABI5 interacts and phosphorylates MPK3. **(A)** Schematic representation of ABI5 domain organization and deletion variants with amino acid numbers used for yeast two hybrid assay. **(B)** Protein – protein interaction analysis of between MPK3 and ABI5 full length and deletion variants by using yeast-two hybrid (Y2H) assay. A, adenine; H, histidine; L, leucine; W, tryptophan. **(C)** BiFC assay showing in-vivo protein– protein interaction between ABI5 and MPK3. The interaction was specifically observed in the nucleus of *N. benthamiana* leaf epidermal cells with YFP signal. No signals were observed with ABI5-nYFP and MPK3-cYFP alone. Arrows indicate the nucleus. Bar = 50μm. **(D)** *In-vitro* phosphorylation assay using bacterially purified ABI5-HIS and GST-MPK3. GST-MPK3 was either incubated alone or with myelin basic protein (MBP) for negative and positive controls, respectively. The reaction mixture was separated on SDS-PAGE and phosphorylation was monitored by autoradiography using typhoon phosphor-imager. **(E)** *In-vitro* phosphorylation assay was performed similar as in **(D)** using ABI5 mutant protein where serine 314 is replaced with alanine (ABI5^S314A^). The upper and lower images represent the autoradiograph and CCB staining, respectively. The plus and minus signs represent the presence and absence of protein in each lane. pMPK3 or pABI5 indicates the phosphorylated protein.

### ABI5 is phosphorylated by MPK3 at serine-314

The *in-vivo* interaction between AtABI5 and AtMPK3 led us to investigate the *in-vitro* phosphorylation assay using bacterially purified AtABI5 and AtMPK3 proteins. We found that AtABI5 is strongly phosphorylated *in-vitro* by AtMPK3 which also showed auto-phosphorylation **(Figure 3D)**. In addition to AtMPK3, AtMPK6 which was activated during ABA treatment also phosphorylated AtABI5 *in-vitro* **(Supplemental Figure 3)**. The phosphorylation of substrates by MAP kinases are mediated at serine/threonine residues followed by a characteristic proline amino acid. Analysis of AtABI5 protein sequence showed a putative serine residue at 314^th^ position **(Supplemental Figure 4A)**. Multiple protein sequence alignment from other plants indicated its evolutionary conservation **(Supplemental Figure 4B)**. We replaced this serine-314 to alanine, a non-phosphorytable amino acid (AtABI5^S314A^) and again performed the *in-vitro* phosphorylation assay using AtMPK3. The phosphorylation of AtABI5^S314A^ was completely abolished **(Figure 3E)** suggesting that MAP kinase phosphorylates AtABI5 at S314 position which is evolutionarily conserved.

### Phosphorylation of AtABI5 by AtMPK3 regulates its subcellular localization and dimerization

To further investigate the role of AtABI5 phosphorylation by MAP kinase *in-vivo*, along with phospho-null variant, AtABI5^S314A^ we generated a phosphor-mimic variant, AtABI5^S314D^ (Serine (S) at 314 was changed to Aspartic acid (D)) and performed localisation of both the variants in tobacco leaves. The localization of phospho-mimic version was solely found in the nucleus **(Figure 4A)**. However, the phospho-null version was localized to the cytosol in addition to the nucleus **(Figure 4B)**. In the cytosol, AtABI5^S314A^ was localised to the peripheral plasma membrane as detected by plasmolysis. Thus, phosphorylation of AtABI5 by MAP kinase might regulate its subcellular localisation. Next, we analysed the dimerization of AtABI5 by BiFC assay using AtABI5 mutant variants. The wild type AtABI5 formed a dimer within the nucleus **(Figure 4C)**. However, interaction between phosphor-null AtABI5 was seen at both cytoplasm and in the nucleus **(Figure 4D)** similar to that of its subcellular localisation. Surprisingly, the interaction was abolished between the phosphor-mimic AtABI5 **(Figure 4E)**. Additionally, no interaction was observed between phosphor-null and phosphor-mimic AtABI5 variants **(Figure 4F)**. No fluorescence was observed in the negative controls **(Supplemental Figure 5)**. Taken together, phosphorylation of AtABI5 not only regulates its subcellular localisation but also inhibits its nuclear dimerization, *in-vivo*.

**Figure 4.**
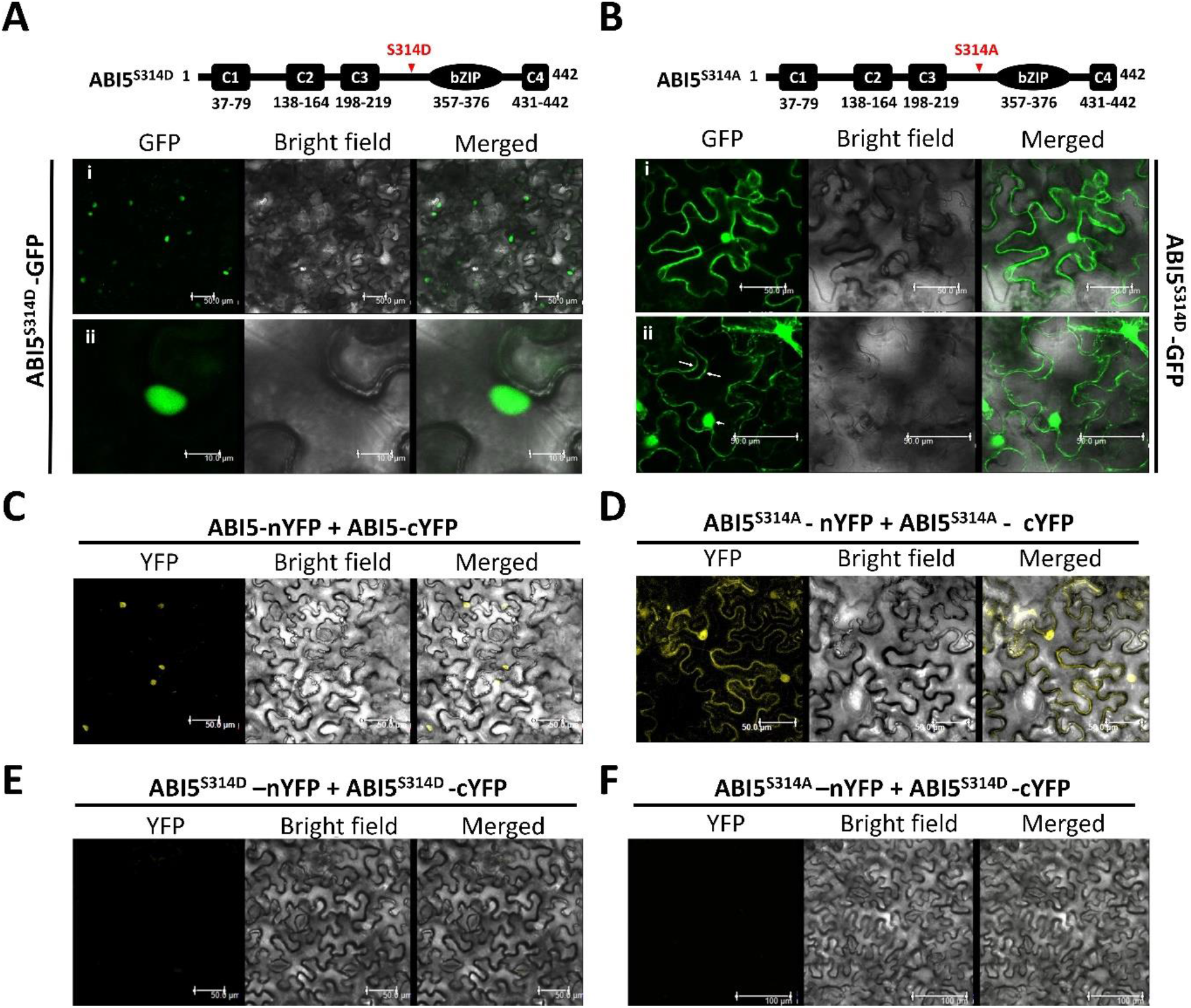
Phosphorylation of ABI5 modulates its subcellular localisation. **(A)** Schematic representation of mutation in ABI5. The upper image represents the phospho-mimic version of ABI5 where serine-314 is replaced with aspartate (ABI5^S314D^). Localization of GFP-ABI5^S314D^ in *N. benthamiana* leaf cells observed under confocal microscope. The GFP fluorescence was detected in the nucleus. **(B)** Schematic representation and localization of GFP-ABI5^S314A^ in *N. benthamiana* leaf cells observed under confocal microscope. Scale bar, 50μm. BiFC analysis of native ABI5 dimerization **(C)**, interaction between AtABI5^S314A^ **(D)**, AtABI5^S314D^ **(E)** and between AtABI5^S314A^-AtABI5^S314D^ **(F)**. Scale bar is indicated in each image.

### MAP kinase mutants are hyposensitive to exogenous ABA during seed germination

ABI5 regulates the seed germination and post-germination growth and positively regulates ABA signaling. Mutation in the AtABI5 leads to ABA insensitive phenotype during seed germination. To examine the role of AtMPK3 during seed germination, we performed ABA sensitivity assay during seed germination in *abi5-8* and *mpk3* mutants along with the wild type Col-0 **(Figure 5A, 5B and 5C)**. As expected we found that wild type seeds were sensitive and *abi5-8* insensitive to exogenous ABA application. Interestingly, *mpk3* mutants exhibited hyposensitive phenotype to ABA during seed germination as compared to wild type. At higher concentrations, the post-germination growth of *mpk3* was arrested as compared to insensitive *abi5-8* mutant. These results suggest that AtMPK3 participates in the ABA inhibition of seed germination, however, hyposensitive phenotype of *mpk3* mutant might be due to the other redundant AtMPK6 which is also activated by ABA and phosphorylates AtABI5.

**Figure 5.**
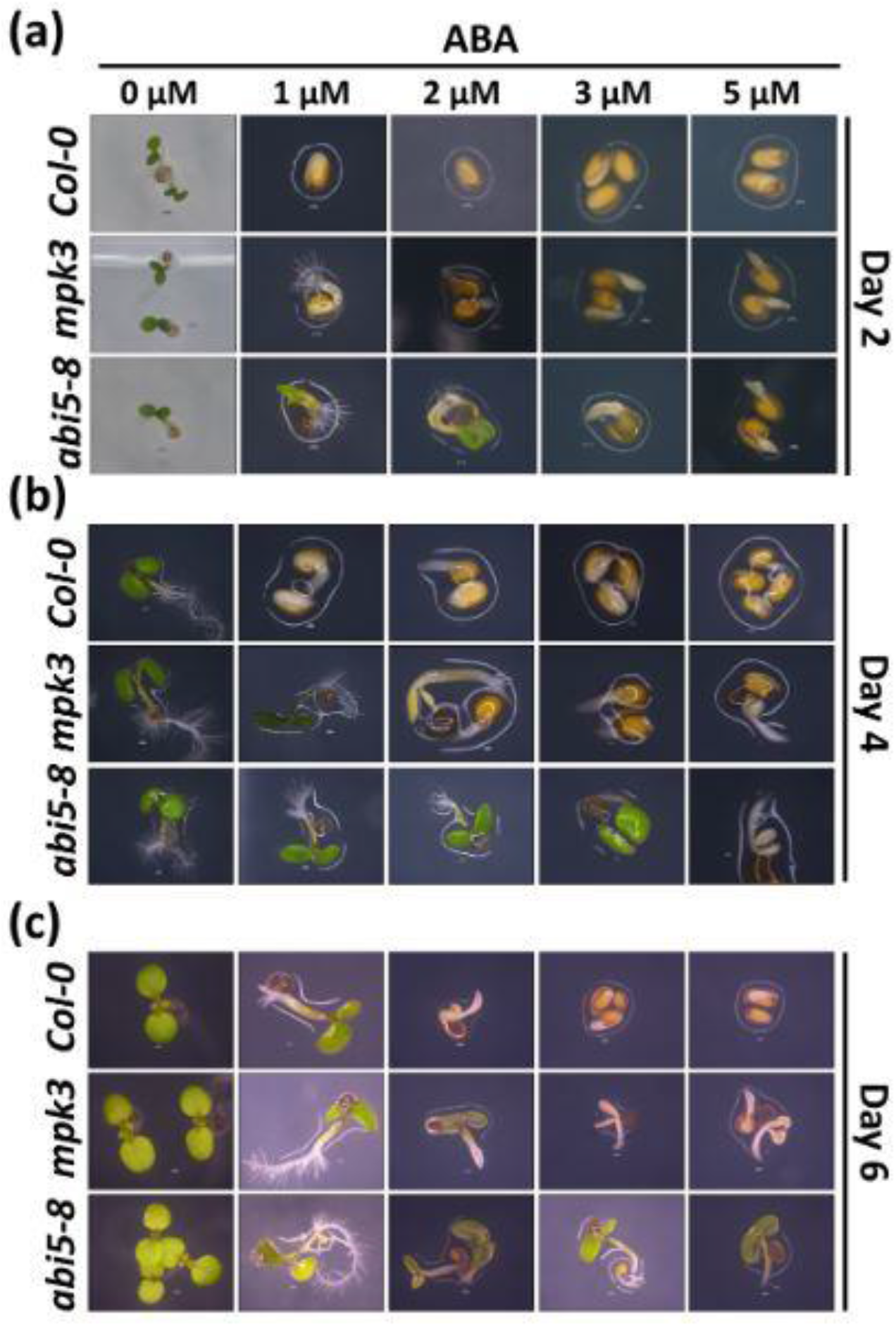
The mpk3 mutant is partially insensitive to exogenous ABA during seed germination. The ABA sensitivity assay was performed by using wild type, *mpk3* and *abi5-8* seeds. Seeds were germinated on different concentrations of ABA (1 μM, 2 μM, 3 μM and 5 μM). Images were captured using Nikon stereomicroscope on 2nd day (a), 3rd day (b) and on 6th day (c) after transferring to light.

### Phosphorylation at serine-314 negatively regulates ABA response during seed germination

To further explain the role of AtABI5 phosphorylation *in-vivo*, we complimented the *abi5-8* mutant by constitutive expression of phosphor-null (AtABI5^S314A^) and phosphor-mimic (AtABI5^S314D^) forms of AtABI5 and performed ABA sensitivity assay. Interestingly, the phosphor-null (AtABI5^S314A^) variant could compliment the *abi5-8* mutant as it showed sensitivity towards exogenous ABA application **(Figure 6A)**. However, the phosphor-mimic AtABI5 showed insensitive phenotype and was unable to compliment the *abi5-8* mutants. These results suggest that phosphorylation of AtABI5 makes it inactive and thus phenotypically mimics the *abi5-8* mutant during seed germination.

**Figure 6.**
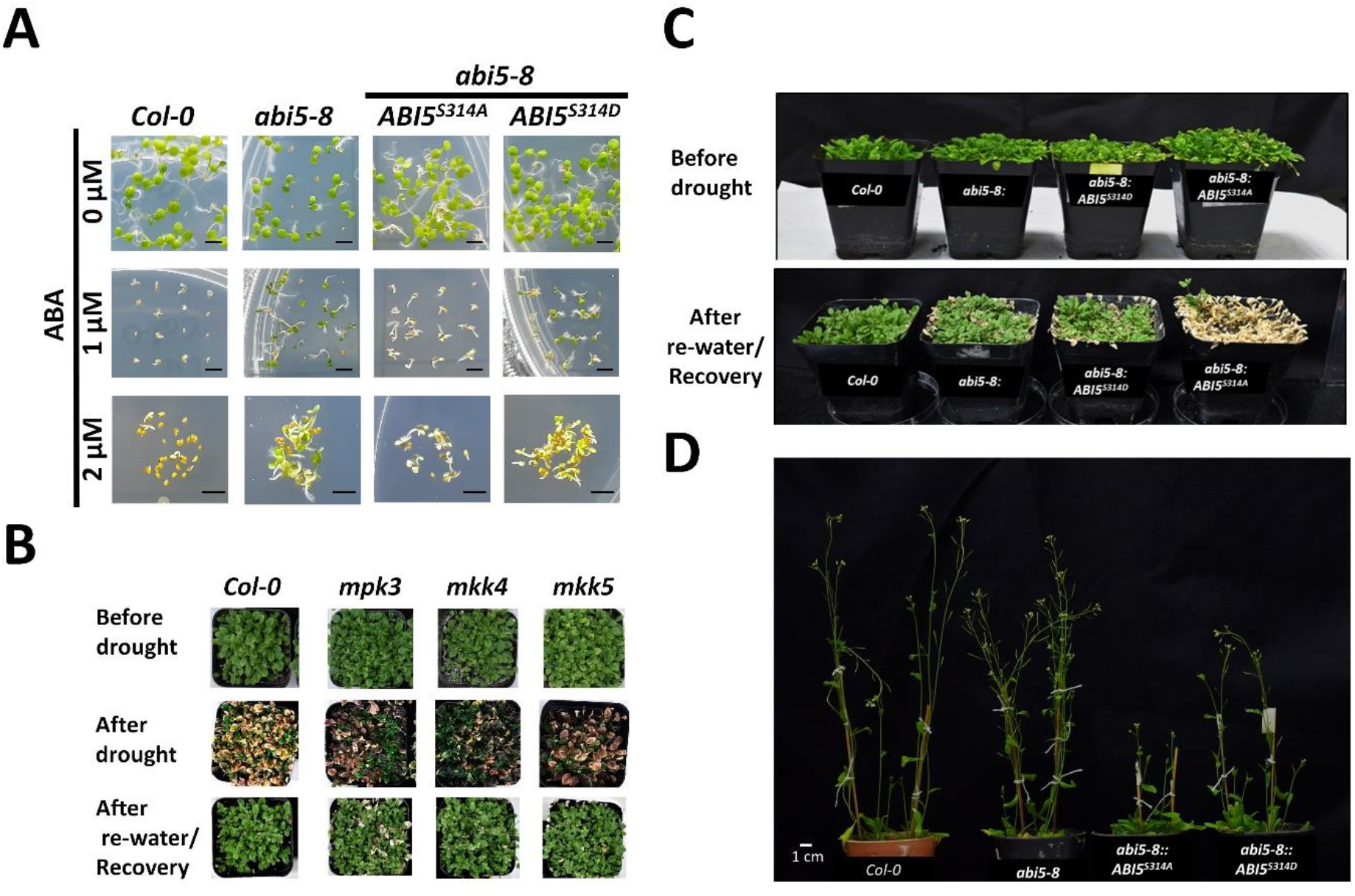
AtMKK4/AtMKK5-AtMPK3/AtMPK6-AtABI5 cascade regulates seed germination, drought stress and floral transition. **(A)** ABA sensitivity assay using wild type, *abi5-8* mutant and ABI5 overexpression lines. The phosphor-null and phosphor-mimic version of ABI5 were constitutively overexpressed in the *abi5-8* background and seed germination were analysed at indicated ABA concentrations. images were taken after 7 days. Scale bar, 1cm. **(B)** Drought stress response of MAP kinase cascade mutants. Phenotypic analysis of wild type, *mpk3*, *mpk6*, *mkk4* and *mkk5* plants subjected water withdrawal and then recovery was monitored after re-watered conditions. **(C)** Phenotype of *abi5-8* and overexpressed phosphor-mimic and phosphor-null ABI5 in *abi5-8* background during drought response. **(D)** Flowering phenotype of ABI5 overexpression and mutants. The flowering response was scored by emergence bolting phenotype.

### Overexpression of phosphor-null ABI5 confers hypersensitive to drought stress

ABA is known to play important role during water deficient conditions or during drought stress. We therefore, were interested to see the effect of ABI5 phosphorylation by MPK3 during drought stress. Towards this end we first analysed the role of MAP kinase component mutants during drought stress. We exposed WT, *mpk3*, *mkk4* and *mkk5* plants to water stress condition by withdrawing water for 14 days before re-watering and assessing the recovery from drought stress. We found that during drought conditions all plants showed chlorosis and leaf damage **(Figure 6B)**. However, MAP kinase mutants, *mpk3*, *mkk4* and *mkk5* showed better tolerance to drought. Even the recovery was faster in MAPK mutants. These results indicate that MAP kinases play a crucial role during drought stress.

Similarly, we further extended our investigation using the *abi5-8* transgenic plants constitutively overexpressing the phospho-null (AtABI5^S314A^) and phospho-mimic (AtABI5^S314D^) forms of AtABI5 during drought stress. We found that *abi5-8* mutant and phospho-mimic AtABI5^S314D^ complimented plants showed insensitive phenotype **(Figure 6C)**. Interestingly, when the mutant is complimented with phosphor-null AtABI5^S314A^ the hypersensitive phenotype to drought was unable to recover during recovery. Taken together, these results clearly suggest that MKK4/MKK5-MPK3-ABI5 module regulates the drought stress response and ABA signaling in Arabidopsis.

### The reversible phosphorylation of ABI5 is crucial for flowering transition

AtABI5 is one of the master regulators of floral transition and reproductive growth. The *abi5* mutants exhibit early flowering phenotype (Wang et al., 2013). To further characterise the role of ABI5 phosphorylation in the flowering response we grew the WT, *abi5-8*, and *abi5-8* mutant lines complimented with AtABI5^S314A^ and AtABI5^S314D^ variants up till flowering **(Figure 6D)**. We found that overexpression of phospho-null AtABI5^S314A^ delayed the flowering as compared to *abi5-8* mutants. However, interestingly it was observed that phospho-mimic variant AtABI5^S314D^ showed intermediate flowering phenotype to that of *abi5-8* and wild type but earlier than that of phosphor-null. The observation indicates that the phosphorylation status of AtABI5 by AtMPK3 also has a role in regulating complex trait like flowering in *A. thaliana*.

## DISCUSSION

ABI5, a bZIP type transcription factor is one of the master regulators which has been well characterized with respect to ABA signaling and stress responses. It has been shown that ABA signaling not only regulates the seed germination but also participates in the floral transition and drought stress where ABI5 is one of the ABA responsive transcription factors (Wang et al., 2013). Phosphorylation of ABI5 by SnRK2 kinases positively regulates the ABA signaling (Nakashima et al., 2009). However, involvement of other unknown kinase in the regulation of this bZIP transcription factor is not so far well characterized. In this study, we report the role of ABA activated MAP kinases, AtMPK3/AtMPK6 in the phospho-regulation of ABI5 during plant development and stress response.

MAP kinase pathways are one of the evolutionary signaling cascades which regulate several developmental and stress responses. MAP kinase cascades are reported to be involved in several ABA signaling including stomatal regulation and stress responses (Wang et al., 2007; Danquah et al., 2014; de Zelicourt et al., 2016). However, there are a very few reports where a direct target of ABA activated MAP kinase has been identified and functionally characterized (Lu et al., 2002). It has been previously shown that an AP2/ERF class transcription factor, ABI4 is phosphorylated by AtMPK3/AtMPK6 and regulates the retrograde signaling in Arabidopsis which is the only well characterised ABA-insensitive transcription factor till now (Guo et al., 2016). In this study, we have identified that ABI5 is an interactor and phosphorylation target of AtMPK3/AtMPK6 by several biochemical and molecular approaches. We found that ABA not only upregulates the AtMPK3 and AtMPK6 kinase activity but selectively induces the transcription and translation of AtMPK3 via AtABI5 transcription factor. In turn, both ABA activated MAP kinases interacts and phosphorylates ABI5.

The promoter analysis of both ABA activated MAP kinases, AtMPK3 and AtMPK6 suggested that only AtMPK3 has the core ABA responsive element (ACGT) while it is absent in AtMPK6 promoter. These observations were further corroborated with the transcript analysis of both kinases where only AtMPK3 showed higher transcript upregulation under ABA treatment. As already reported, we also found that under ABA treatment the expression of AtABI5 was upregulated (Liu and Stone, 2010). To find the role of AtABI5 in the transcriptional regulation of AtMPK3, we performed immunoblot analysis of AtMPK3 protein in *abi5-8* mutant and the data clearly indicated lower accumulation of its protein under ABA treatment. However, the involvement of other ABA-responsive TFs along with AtABI5 where AtMPK3 protein was reduced as compared to wild type but not completely abolished cannot be ruled out. The binding of AtABI5 to the promoter of AtMPK3 was confirmed by EMSA using bacterially purified AtABI5 protein and AtMPK3 promoter fragment.

The feedback regulation in several signaling pathways have been reported where the TF regulating the expression of MAPKs were also the targets of respective MAPKs (Sethi et al., 2014; Singh and Sinha, 2016). In the similar line, we found that AtABI5 along with transcriptionally upregulating AtMPK3 also shows protein-protein interaction with it. The *in-vivo* interaction was observed in the nucleus where AtABI5 is known to regulate the transcripts of ABA-responsive genes (Liu and Stone, 2013). The AtABI5 protein has been shown to undergo through multiple post-translational modifications including phosphorylation by SnRKs and BIN2 including ubiquitination/SUMOylation **(Supplemental Figure 6)** (Miura et al., 2009; Nakashima et al., 2009; Liu and Stone, 2010, 2014). The *in-vitro* phosphorylation assay and site-directed mutagenesis conferred that AtABI5 is phosphorylated also by AtMPK3 and that this phosphorylation is at evolutionarily conserved serine residue present at 314^th^ position. We also found that other ABA-activated AtMPK6 which is also redundant in function can phosphorylate AtABI5. Several reports suggest that protein activity and subcellular localization of most proteins and AtABI5 are regulated by post translational modifications (Liu and Stone, 2010, 2013; Guo et al., 2016). Our subcellular localization studies using phosphor-null and phosphor-mimic variant, AtABI5^S314A^ and AtABI5^S314D^, respectively showed a contrasting phenotype. The phosphor-mimic AtABI5^S314D^ was exclusively localised to the nucleus, as the wild type AtABI5 reported before (Liu and Stone, 2013). However, the phosphor-null AtABI5 was localised also in the cytoplasm along with the nucleus. Thus phosphorylation of AtABI5 by MAP kinase at Serine-314 may regulate its subcellular localization and possibly its cytoplasmic stability. This can be true as mutation in the putative ubiquitination site in AtABI5 or use of MG132 inhibitor of proteasome significantly enhanced the wild type protein in the cytoplasm (Liu and Stone, 2010, 2013) which is in corroboration with our observation of ABI5^S314A^ localization. Next, we also found that AtABI5 phosphorylation inhibits the homodimer formation in the nucleus (Nakamura et al., 2001). Therefore, it seems like both phosphorylation and dephosphorylation of AtABI5 is essential for its subcellular activity (Dai et al., 2013).

Next we functionally characterised the role of AtABI5 phosphorylation by MAP kinases *in-vivo*. We observed that overexpression of phospho-null AtABI5^S314A^ in *abi5-8* mutant background restored the ABA sensitivity to that of *Col-0* during seed germination under ABA treatment. Meanwhile, overexpression of phosphor-mimic AtABI5^S314D^ failed to rescue the phenotype and exhibited the phenotype of *abi5-8* mutant. After functionally characterising the role of phosphorylation during seed germination, we investigated the developmental role of AtABI5 during floral transition. Previously, it has been shown that ABA delays the floral transition through AtABI5 which binds to the promoter of *FLOWERING LOCUS C* (*FLC*) (Wang et al., 2013). They reported that *abi5* mutants show early flowering and overexpression significantly delays the same. Our finding suggested that both mutant ABI5 versions delayed the flowering but phosphor-null AtABI5 showed severe delay as compared to phosphor-mimic form which showed intermediate flowering phenotype. Thus we concluded that reversible phosphorylation of AtABI5 by AtMPK3 is crucial for floral transition.

ABA signaling is also activated during water deficient condition or drought stress and MAP kinase cascade are also known to regulate the stress response. Thus we further investigated the role of MAP kinase-ABI5 module during drought stress and recovery. Our data suggest that *mpk3, mkk4* and *mkk5* showed drought stress tolerance. We further found that *abi5-8* mutant and phospho-mimic AtABI5^S314D^ transgenic plants also showed similar drought insensitive phenotype as MAP kinase mutants. However, phosphor-null AtABI5^S314A^ was hypertensive to drought. These data suggest that MAP kinase and AtABI5 regulate the drought stress response in an ABA-dependent manner.

In summary, we can conclude that ABA regulates the activity of AtMPK3/AtMPK6 downstream of AtMKK4/AtMKK5. Moreover, AtMPK3 expression is also subjected to transcriptional regulation downstream of AtABI5. The activated AtMPK3 and AtMPK6, in-turn phosphorylate AtABI5 at serine-314 position which regulates its subcellular localization and dimerization and interacts with AtMPK3 promoter in a positive feedback mechanism. This reversible phosphorylation functionally regulates the AtABI5 during seed germination, floral transition and drought stress in Arabidopsis. A simple working model of the involvement of MAPK in ABI5 function is depicted in **Figure 7**. It is likely that MAP kinase cascade and ABA signaling crosstalk at multiple developmental pathways and during stress response and this study provides an extensive effort towards understanding this molecular mechanism.

**Figure 7.**
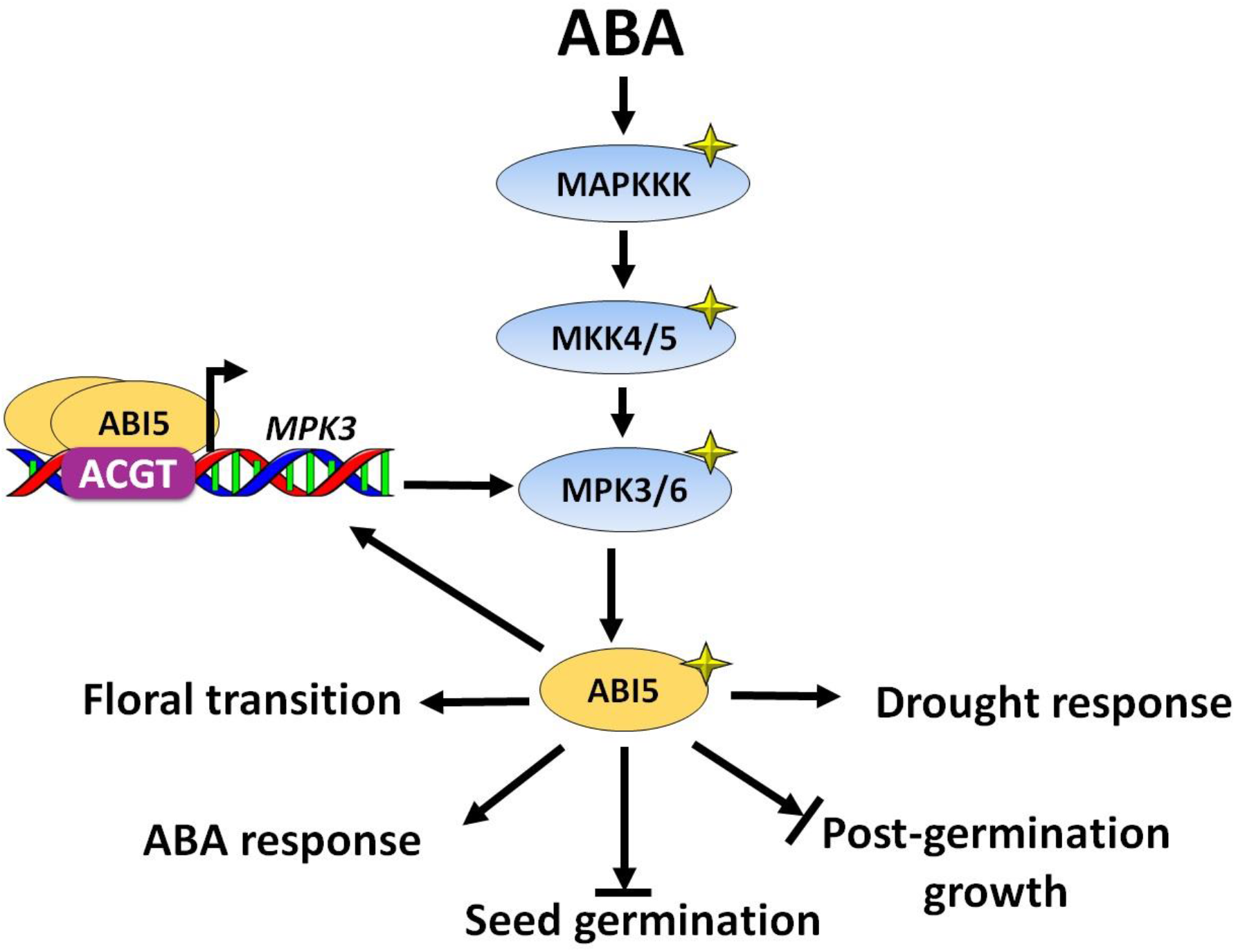
A proposed model showing the role of MAP kinases cascade upstream of ABI5 during ABA signaling. Increased ABA due to stress of developmental cues may results into the activation of MAP kinase cascade where MKK4/5 is activated by upstream unknown MAKKK. The MPK3 and MPK6 are activated by MKK4/5 that interact and phosphorylate ABI5 that regulates diverse physiological response such as inhibition of seed germination and post-germination growth, drought stress and floral transition. In turn ABI5 selectively interacts with AtMPK3 promoter and upregulates its transcript as well as protein level in an ABA-dependent manner. The open and closed arrow heads indicate the promotion and inhibition, respectively.

## Accession Numbers

The nucleotide and protein sequences used in this article can be found in the TAIR database under following accession numbers: AT3G45640 (MPK3), AT2G43790 (MPK6), AT2G36270 (ABI5), AT1G51660 (MKK4), AT3G21220 (MKK5), AT3G18780 (ACTIN2). The *mpk3* (SALK_151594), *mpk6* (Salk_073907) was used from our previous study and the following *abi5-8* line (SALK_013163), *mkk4* (SALK_058307), *mkk5-1* (SALK_047797) were obtained from the Arabidopsis Biological Resource Center.

## Acknowledgements

This work was supported by core grants from the National Institute of Plant Genome Research from the Department of Biotechnology (DBT), Government of India. PKB and DV are recipients of fellowship from the DBT, University Grant Commission (UGC) Government of India, respectively. NV is recipient of fellowship from DBT-Biocare Women Scientist scheme. Authors thank the Radioisotope facility, Confocal Microscopy Facility and the Central Instrumentation Facility of NIPGR, New Delhi, India. The authors are thankful to DBT-eLibrary Consortium (DeLCON) for providing access to e-resources.

## Author Contributions

PKB and AKS designed the experiments and overall study. PKB performed all the extensive experimental work with the help of DV and NV. PKB wrote the first draft of the manuscript. All participated in the data preparation and AKS participated in the discussion and progress of work. Final draft was approved by AKS.

## Supplemental Data

**Supplemental Figure 1:**
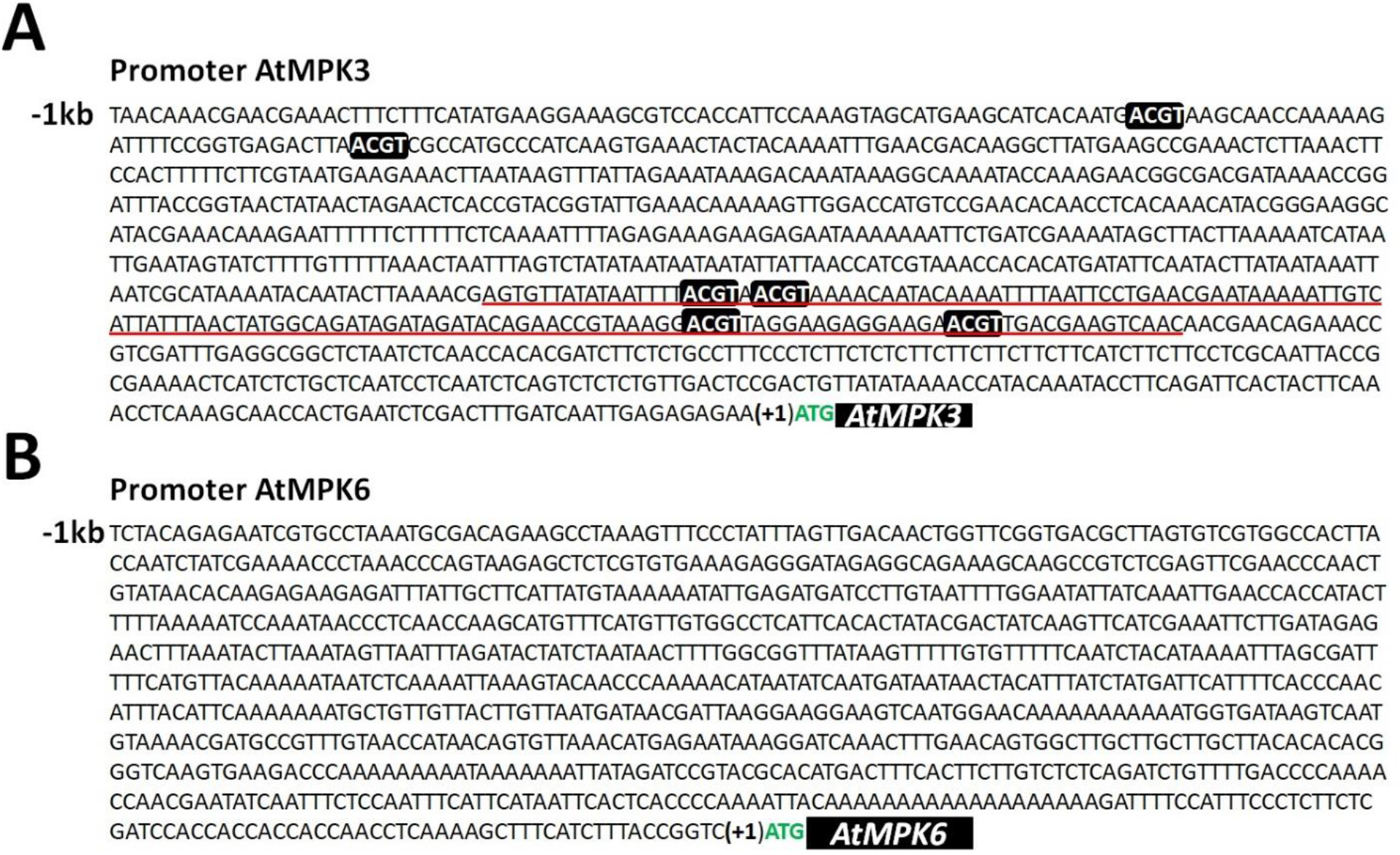
Promoter sequence of MPK3 and MPK6. DNA sequence showing the upstream 1kb promoter elements from ATG of **(A)** AtMPK3 and **(B)** AtMPK6. The core ABREs (ACGT) are in black boxes. Red underline DNA sequence was used for EMSA.

**Supplemental Figure 2:**
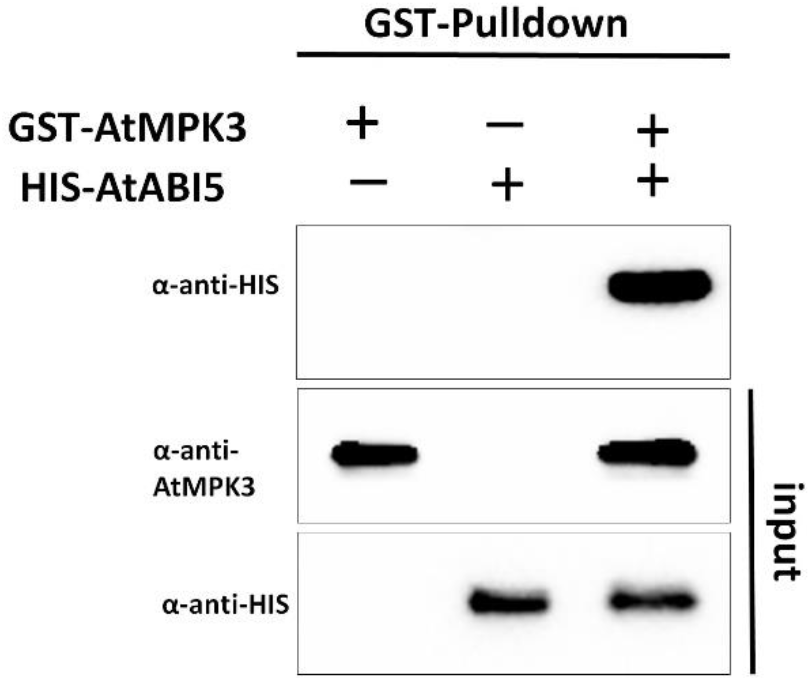
Pull down assay of MPK3 and ABI5 interaction. *In-vitro* GST-pulldown assay showing the interaction between GST-AtMPK3 and HIS-AtABI5. Bacterially purified GST-AtMPK3 and His-AtABI5 proteins were pulldown using GST-sepharose beads and detected by Anti-HIS and Anti-AtMPK3 antibody. Plus (+) and minus (−) sings indicate the presence and absence of proteins, respectively.

**Supplemental Figure 3:**
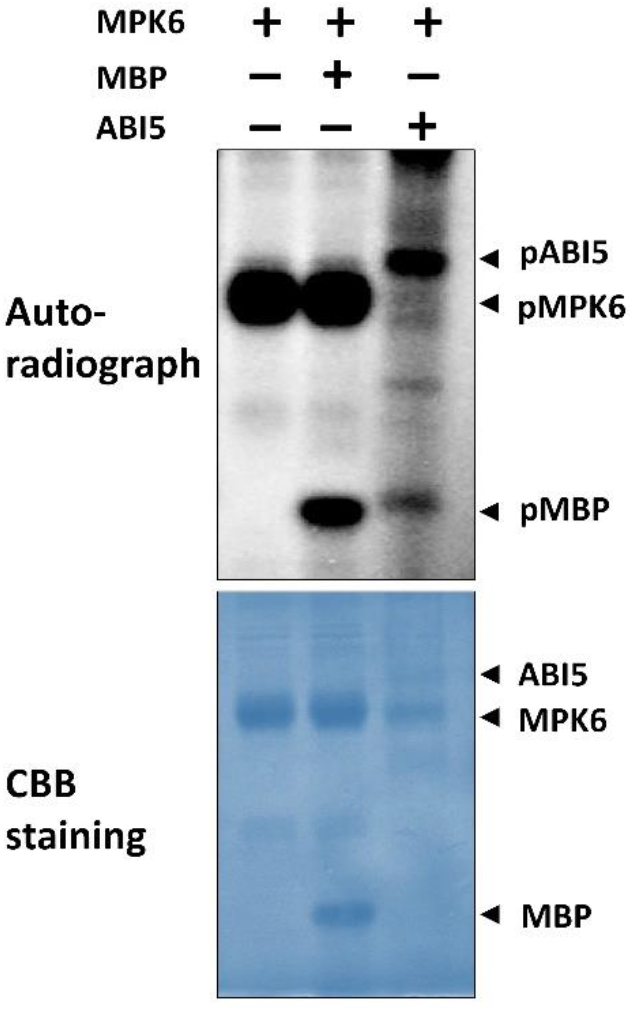
Phosphorylation of ABI5 by MPK6. *In-vitro* phosphorylation assay showing the phosphorylation of ABI5 by AtMPK6. Recombinant His-AtMPK6 was either incubated alone or with MBP and His-ABI5 in kinase buffer and phosphorylation of ABI5 was detected by auto-radiography. Plus (+) and minus (−) sings represent the presence and absence of proteins in each lane.

**Supplemental Figure 4:**
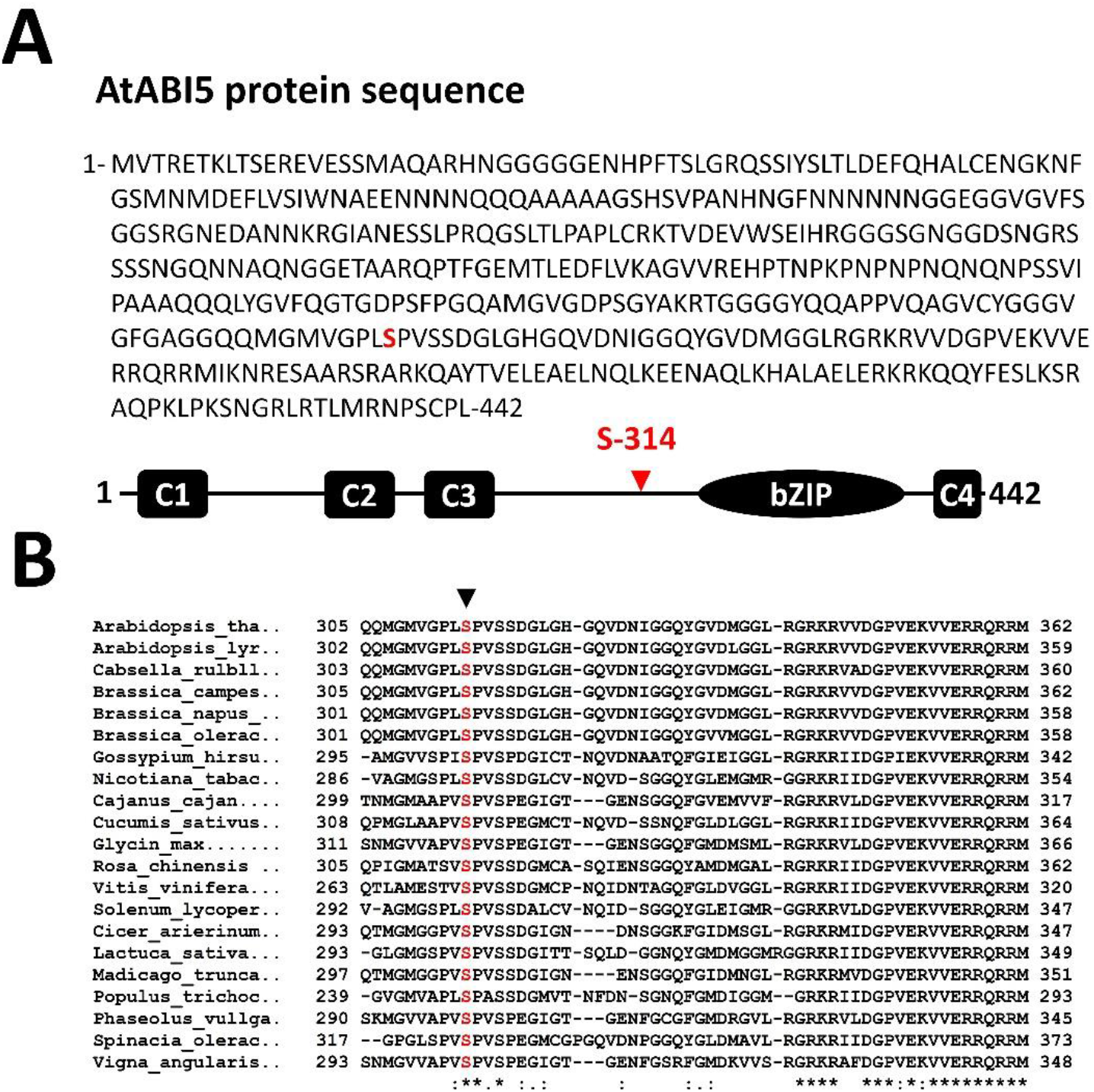
MAP kinase phosphorylation sites on ABI5. **(A)** Protein sequence of ABI5 showing the putative serine residue as a MAP kinase phosphorylation site (Red). **(B)** Multiple protein sequence alignment showing the evolutionary conservation of seine-314 of AtABI5 in other plants. Arrow head indicating the invariant serine residue.

**Supplemental Figure 5:**
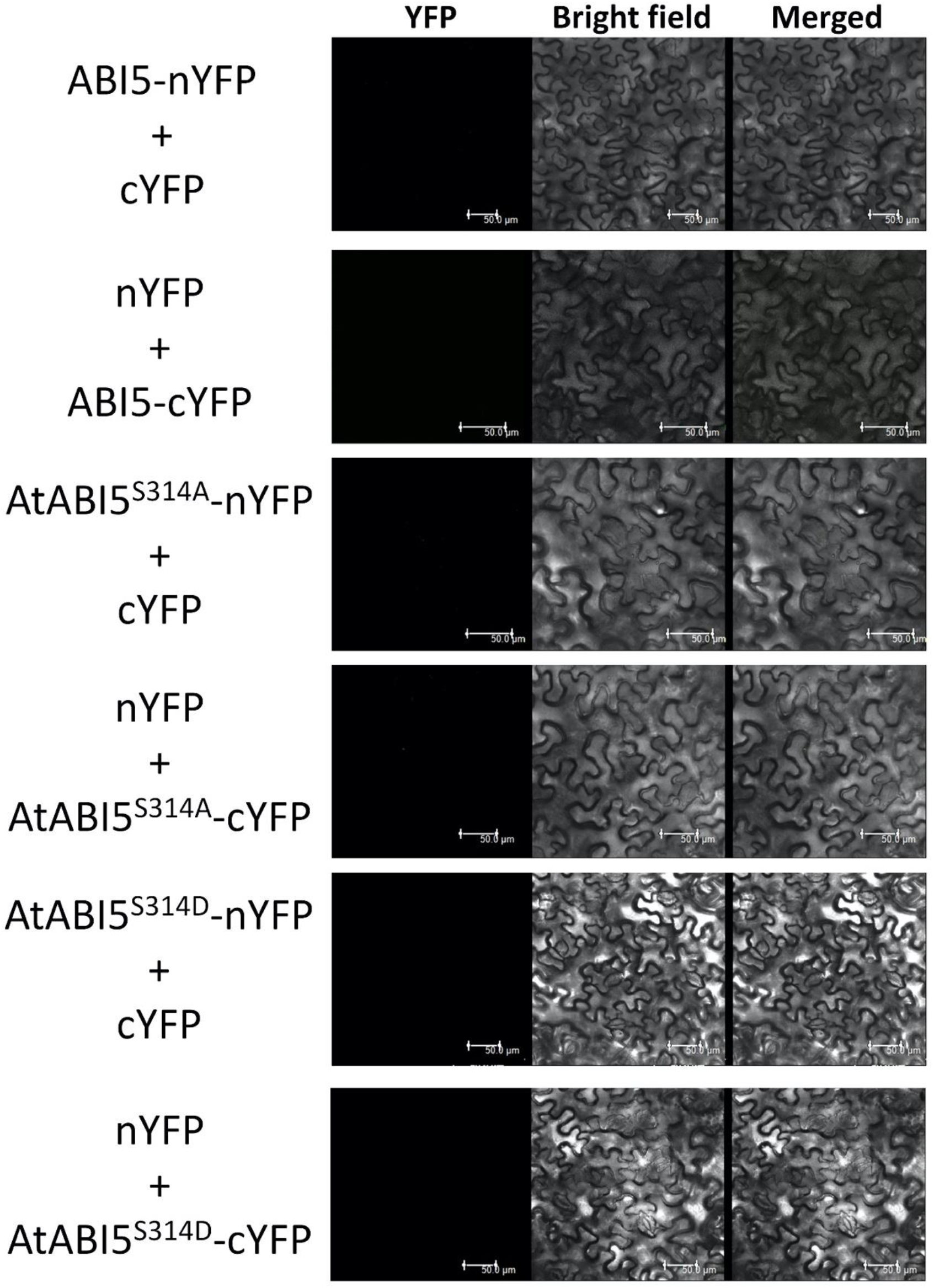
Controls for BiFC assay. BiFC negative controls of AtABI5-AtMPK3 protein-protein interaction in *N. benthamiana* leaves (Related to Figure 4C-4F). Bar = 50μm.

**Supplemental Figure 6:**
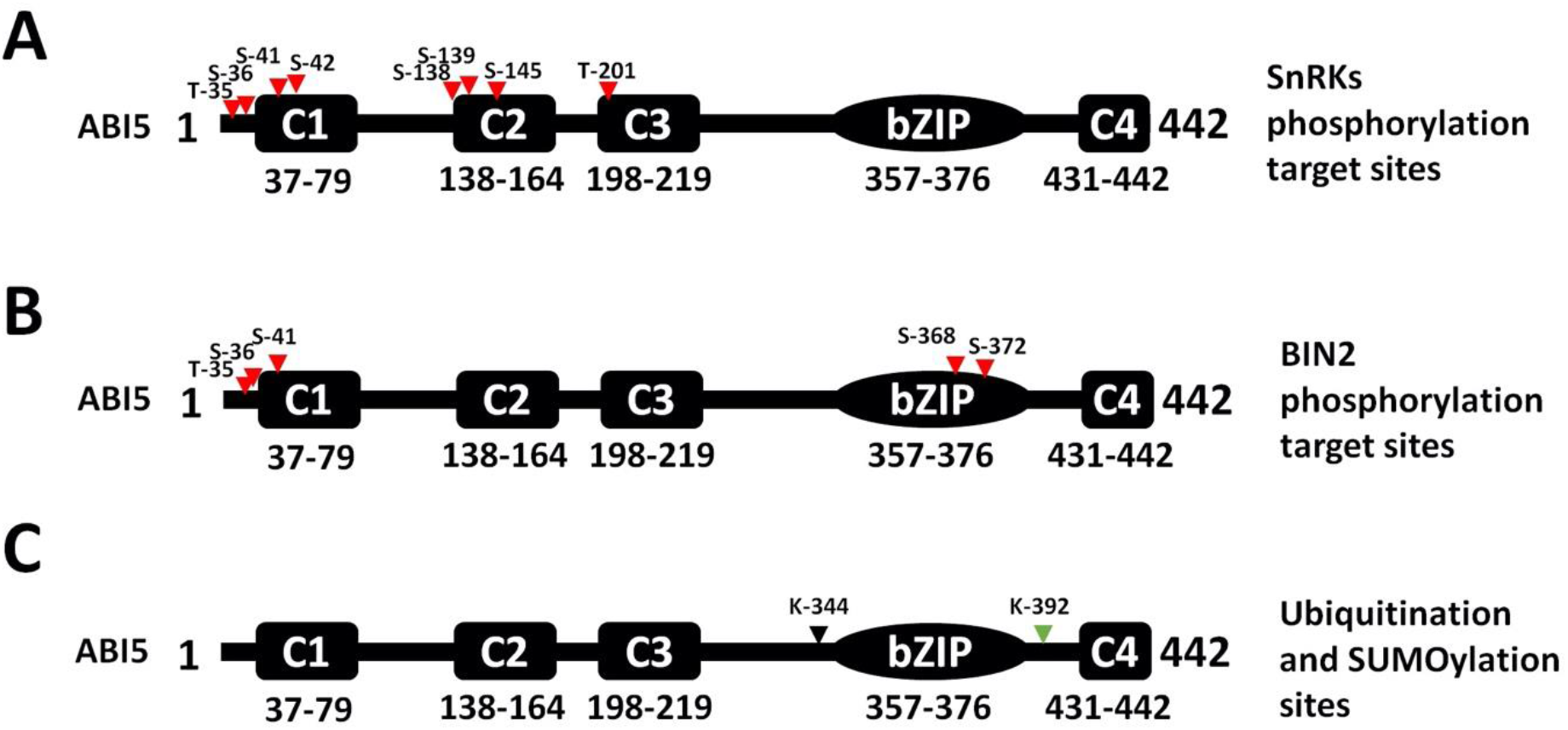
Schematic representation of post-translational modification of ABI5. **(A)** SnRK2 specific phosphorylation sites on ABI5 protein. **(B)** BIN2 regulated phosphorylation sites on ABI5. **(C)** Ubiquitinated and SUMOylated lysine residues by KEG and SIZ1. Red, black and green triangles represent phosphorylation, ubiquitination and SUMOylation sites on ABI5.

**Supplemental Table 1:**
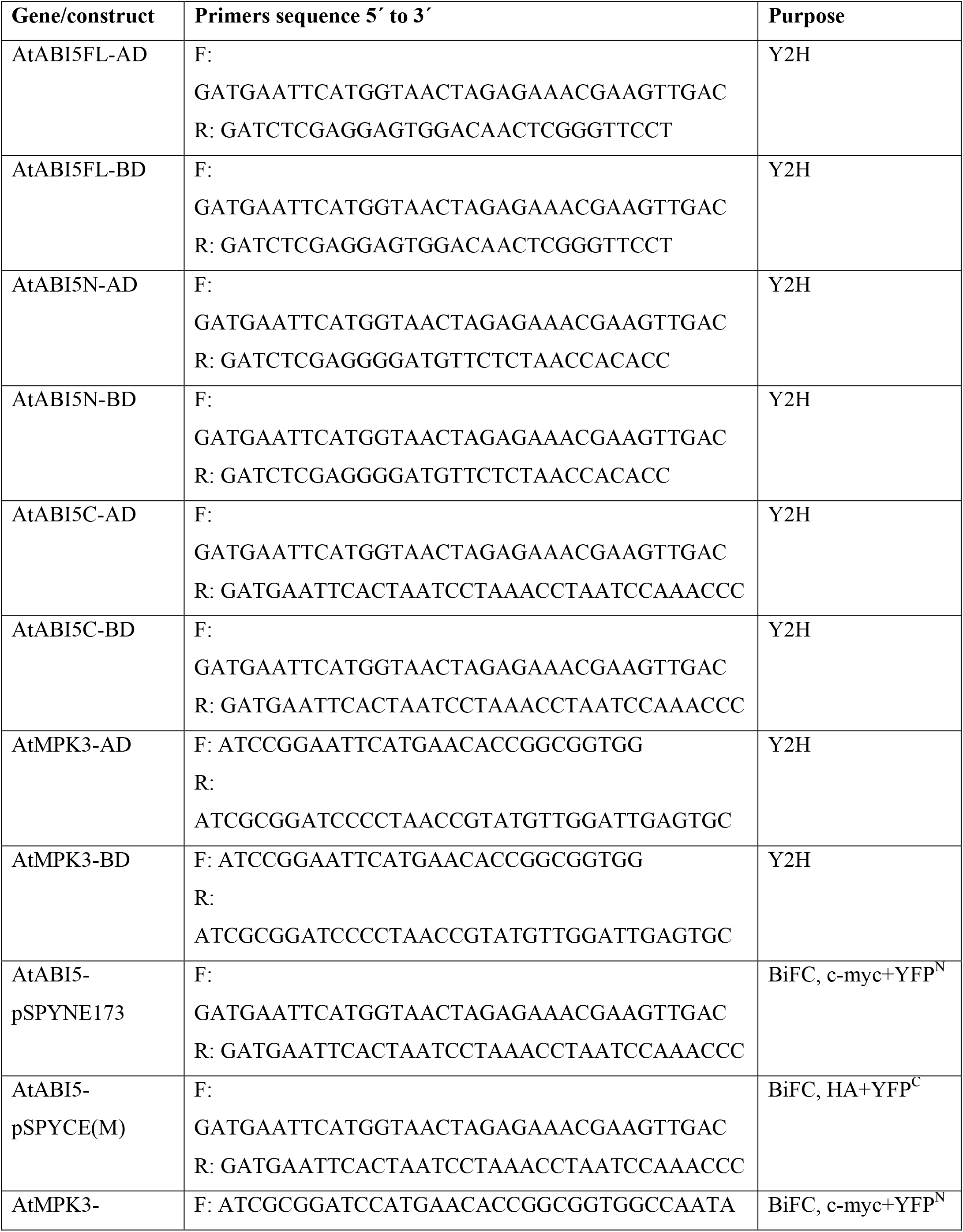
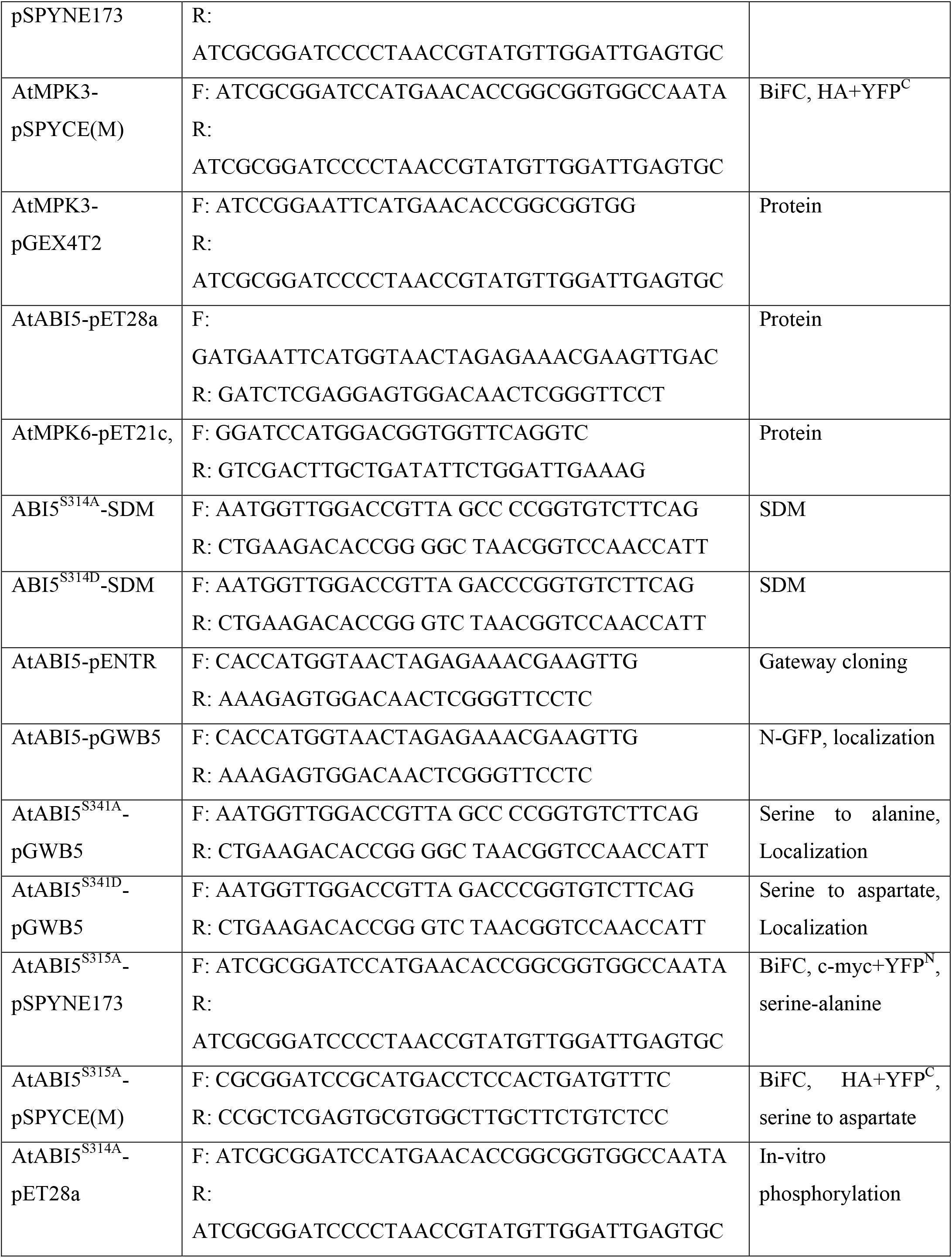
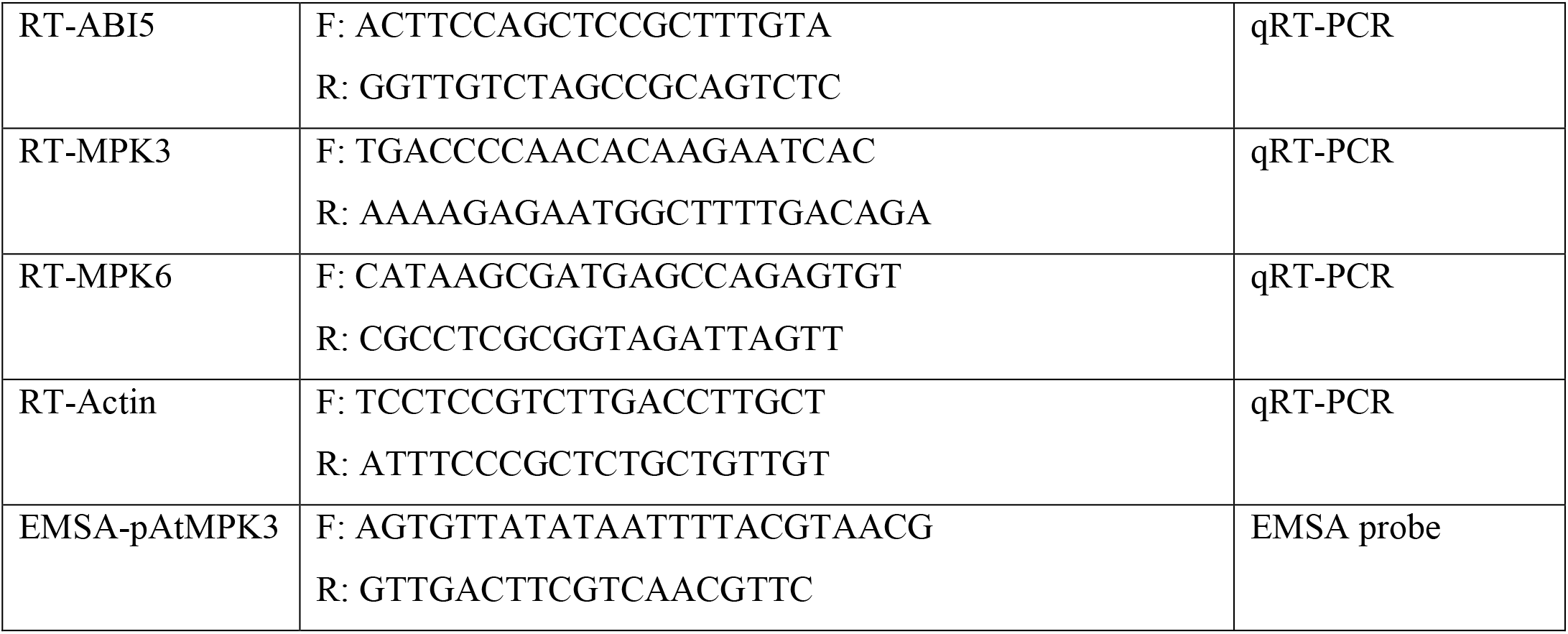
Primer list used in the study.

## References

Dai, M.Q., Xue, Q., McCray, T., Margavage, K., Chen, F., Lee, J.H., Nezames, C.D., Guo, L.Q., Terzaghi, W., Wan, J.M., Deng, X.W., and Wang, H.Y. (2013). The PP6 Phosphatase Regulates ABI5 Phosphorylation and Abscisic Acid Signaling in Arabidopsis. Plant Cell 25, 517–534.

Danquah, A., de Zelicourt, A., Colcombet, J., and Hirt, H. (2014). The role of ABA and MAPK signaling pathways in plant abiotic stress responses. Biotechnology Advances 32, 40–52.

Danquah, A., de Zelicourt, A., Boudsocq, M., Neubauer, J., Frey, N.F.D., Leonhardt, N., Pateyron, S., Gwinner, F., Tamby, J.P., Ortiz-Masia, D., Marcote, M.J., Hirt, H., and Colcombet, J. (2015). Identification and characterization of an ABA-activated MAP kinase cascade in Arabidopsis thaliana. Plant Journal 82, 232–244.

de Zelicourt, A., Colcombet, J., and Hirt, H. (2016). The Role of MAPK Modules and ABA during Abiotic Stress Signaling. Trends in Plant Science 21, 677–685.

Fujii, H., Verslues, P.E., and Zhu, J.-K. (2007). Identification of two protein kinases required for abscisic acid regulation of seed germination, root growth, and gene expression in Arabidopsis. Plant Cell 19, 485–494.

Guo, H., Feng, P., Chi, W., Sun, X., Xu, X., Li, Y., Ren, D., Lu, C., Rochaix, J.D., Leister, D., and Zhang, L. (2016). Plastid-nucleus communication involves calcium-modulated MAPK signalling. Nature Communications 7.

Harb, A., Krishnan, A., Ambavaram, M.M.R., and Pereira, A. (2010). Molecular and Physiological Analysis of Drought Stress in Arabidopsis Reveals Early Responses Leading to Acclimation in Plant Growth. Plant Physiology 154, 1254–1271.

Jalmi, S.K., Bhagat, P.K., Verma, D., Noryang, S., Tayyeba, S., Singh, K., Sharma, D., and Sinha, A.K. (2018). Traversing the Links between Heavy Metal Stress and Plant Signaling. Frontiers in Plant Science 9.

Jammes, F., Song, C., Shin, D.J., Munemasa, S., Takeda, K., Gu, D., Cho, D., Lee, S., Giordo, R., Sritubtim, S., Leonhardt, N., Ellis, B.E., Murata, Y., and Kwak, J.M. (2009). MAP kinases MPK9 and MPK12 are preferentially expressed in guard cells and positively regulate ROS-mediated ABA signaling. Proceedings of the National Academy of Sciences of the United States of America 106, 20520–20525.

Khokon, M.A.R., Salam, M.A., Jammes, F., Ye, W., Hossain, M.A., Uraji, M., Nakamura, Y., Mori, I.C., Kwak, J.M., and Murata, Y. (2015). Two guard cell mitogen-activated protein kinases, MPK9 and MPK12, function in methyl jasmonate-induced stomatal closure in Arabidopsis thaliana. Plant Biology 17, 946–952.

Li, Q., Wang, Y.-J., Liu, C.-K., Pei, Z.-M., and Shi, W.-L. (2017a). The crosstalk between ABA, nitric oxide, hydrogen peroxide, and calcium in stomatal closing of Arabidopsis thaliana. Biologia 72, 1140–1146.

Li, Y., Cai, H., Liu, P., Wang, C., Gao, H., Wu, C., Yan, K., Zhang, S., Huang, J., and Zheng, C. (2017b). Arabidopsis MAPKKK18 positively regulates drought stress resistance via downstream MAPKK3. Biochemical and Biophysical Research Communications 484, 292–297.

Liu, H., and Stone, S.L. (2010). Abscisic Acid Increases Arabidopsis ABI5 Transcription Factor Levels by Promoting KEG E3 Ligase Self-Ubiquitination and Proteasomal Degradation. Plant Cell 22, 2630–2641.

Liu, H., and Stone, S.L. (2013). Cytoplasmic Degradation of the Arabidopsis Transcription Factor ABSCISIC ACID INSENSITIVE 5 Is Mediated by the RING-type E3 Ligase KEEP ON GOING. Journal of Biological Chemistry 288, 20267–20279.

Liu, H., and Stone, S.L. (2014). Regulation of ABI5 turnover by reversible post-translational modifications. Plant Signaling & Behavior 9.

Lopez-Molina, L., and Chua, N.H. (2000). A null mutation in a bZIP factor confers ABA-insensitivity in Arabidopsis thaliana. Plant and Cell Physiology 41, 541–547.

Lopez-Molina, L., Mongrand, S., and Chua, N.H. (2001). A postgermination developmental arrest checkpoint is mediated by abscisic acid and requires the AB15 transcription factor in Arabidopsis. Proceedings of the National Academy of Sciences of the United States of America 98, 4782–4787.

Lopez-Molina, L., Mongrand, B., McLachlin, D.T., Chait, B.T., and Chua, N.H. (2002). ABI5 acts downstream of ABI3 to execute an ABA-dependent growth arrest during germination. Plant Journal 32, 317–328.

Lu, C., Han, M.H., Guevara-Garcia, A., and Fedoroff, N.V. (2002). Mitogen-activated protein kinase signaling in postgermination arrest of development by abscisic acid. Proceedings of the National Academy of Sciences of the United States of America 99, 15812–15817.

Matsuoka, D., Yasufuku, T., Furuya, T., and Nanmori, T. (2015). An abscisic acid inducible Arabidopsis MAPKKK, MAPKKK18 regulates leaf senescence via its kinase activity. Plant Molecular Biology 87, 565–575.

Miura, K., Lee, J., Jin, J.B., Yoo, C.Y., Miura, T., and Hasegawa, P.M. (2009). Sumoylation of ABI5 by the Arabidopsis SUMO E3 ligase SIZ1 negatively regulates abscisic acid signaling. Proceedings of the National Academy of Sciences of the United States of America 106, 5418–5423.

Nakamura, S., Lynch, T.J., and Finkelstein, R.R. (2001). Physical interactions between ABA response loci of Arabidopsis. Plant Journal 26, 627–635.

Nakashima, K., Fujita, Y., Kanamori, N., Katagiri, T., Umezawa, T., Kidokoro, S., Maruyama, K., Yoshida, T., Ishiyama, K., Kobayashi, M., Shinozaki, K., and Yamaguchi-Shinozaki, K. (2009). Three Arabidopsis SnRK2 Protein Kinases, SRK2D/SnRK2.2, SRK2E/SnRK2.6/OST1 and SRK2I/SnRK2.3, Involved in ABA Signaling are Essential for the Control of Seed Development and Dormancy. Plant and Cell Physiology 50, 1345–1363.

Piskurewicz, U., Jikumaru, Y., Kinoshita, N., Nambara, E., Kamiya, Y., and Lopez-Molina, L. (2008). The Gibberellic Acid Signaling Repressor RGL2 Inhibits Arabidopsis Seed Germination by Stimulating Abscisic Acid Synthesis and ABI5 Activity. Plant Cell 20, 2729–2745.

Raghuram, B., Sheikh, A.H., Rustagi, Y., and Sinha, A.K. (2015). MicroRNA biogenesis factor DRB1 is a phosphorylation target of mitogen activated protein kinase MPK3 in both rice and Arabidopsis. Febs Journal 282, 521–536.

Sethi, V., Raghuram, B., Sinha, A.K., and Chattopadhyay, S. (2014). A Mitogen-Activated Protein Kinase Cascade Module, MKK3-MPK6 and MYC2, Is Involved in Blue Light-Mediated Seedling Development in Arabidopsis. Plant Cell 26, 3343–3357.

Sheikh, A.H., Raghuram, B., Jalmi, S.K., Wankhede, D.P., Singh, P., and Sinha, A.K. (2013). Interaction between two rice mitogen activated protein kinases and its possible role in plant defense. Bmc Plant Biology 13.

Shu, K., Chen, F., Zhou, W.G., Luo, X.F., Dai, Y.J., Shuai, H.W., and Yang, W.Y. (2018). ABI4 regulates the floral transition independently of ABI5 and ABI3. Molecular Biology Reports 45, 2727–2731.

Singh, P., and Sinha, A.K. (2016). A Positive Feedback Loop Governed by SUB1A1 Interaction with MITOGEN-ACTIVATED PROTEIN KINASE3 Imparts Submergence Tolerance in Rice. Plant Cell 28, 1127–1143.

Verma, D., Jalmi, S.K., Bhagat, P.K., Verma, N., and Sinha, A.K. (2019). A bHLH transcription factor, MYC2, imparts salt intolerance by regulating proline biosynthesis in Arabidopsis. The FEBS Journal 129, 238–243.

Wang, H., Ngwenyama, N., Liu, Y., Walker, J.C., and Zhang, S. (2007). Stomatal development and patterning are regulated by environmentally responsive mitogen-activated protein kinases in Arabidopsis. Plant Cell 19, 63–73.

Wang, Y.P., Li, L., Ye, T.T., Lu, Y.M., Chen, X., and Wu, Y. (2013). The inhibitory effect of ABA on floral transition is mediated by ABI5 in Arabidopsis. Journal of Experimental Botany 64, 675–684.

Xu, J., and Zhang, S. (2015). Mitogen-activated protein kinase cascades in signaling plant growth and development. Trends in Plant Science 20, 56–64.

